# The zebra finch auditory cortex reconstructs occluded syllables in conspecific song

**DOI:** 10.1101/2021.07.19.452925

**Authors:** Bao Le, Margot C Bjoring, C Daniel Meliza

**Affiliations:** Department of Psychology, University of Virginia, Charlottesville VA 22901, USA; Program in Fundamental Neuroscience, University of Virginia, Charlottesville VA 22901, USA

## Abstract

In the perceptual illusion known as phonemic (or auditory) restoration, listeners hear sounds occluded by short bursts of noise. The neural mechanisms that create this illusion by generating predictions of the missing information remain poorly understood. Zebra finches (*Taeniopygia guttata*) use song, a sequence of complex vocal elements, to communicate in noisy social environments. Here, we found that in anesthetized finches, populations of single units in the homolog of auditory cortex respond to occluded songs as if the missing elements were present. This occurs even for songs birds have never heard, but not if the context is masked or lacks species-typical syntax. These results suggest that local neural dynamics pre-attentively instantiate a general model of conspecific song that biases auditory responses to restore missing information.

## Introduction

Sensory inputs are often incomplete and degraded by noise. To compensate, the brain uses internal models constructed from context, experience, and innate biases ^1 –4^. Perceptual illusions occur when these models diverge from reality ^5^, as in the auditory illusion of phonemic restoration. When speech phonemes are deleted and replaced with noise, listeners hear the missing phonemes behind the occluding noise ^6^,^7^. The restored phoneme is biased by phonological and lexical context ^8^,^9^ as well as the listener’s native language ^10^,^11^, consistent with an internal model shaped by experience that uses context to create illusory activity corresponding to missing auditory input. The neural mechanisms underlying internal sensory models remain poorly understood. Auditory restoration is thought to involve an interaction between bottom-up biases and model-based expectations ^4^,^9^,^12^. Studies in nonhuman animals have only examined neural responses using simple tones ^13^,^14^, which lack the complex temporal structure to support model-based contextual prediction necessary for restoring occluded vocalizations ^15 –17^.

In this study, we examined auditory restoration in the zebra finch, a songbird with an acoustically and syntactically rich song ^18^ used in noisy social settings where an internal model would benefit receptive communication. Zebra finch songs are sequences of vocal elements (“syllables”) that are highly stereotyped within an individual’s repertoire ^19^, and neurons in the auditory pallium, which is homologous to the mammalian auditory cortex ^20^, are selective for specific syllables ^21^,^22^. We hypothesized that experience hearing zebra finch songs strengthens the synaptic connections between neurons tuned to successive syllables, creating a manifold that biases population dynamics towards trajectories corresponding to the next vocal element in familiar songs, even in the absence of a physical syllable (Fig. 1A). Thus, without excluding a potential role for top-down modulation from higher-order areas, we expect to find local biases toward familiar songs in the auditory pallium in the absence of attention. To test this idea, we recorded from the auditory pallium in anesthetized zebra finches, predicting that responses to occluded songs would resemble responses to stimuli corresponding to the illusory percept (i.e., illusory responses) and not to the stimuli that were physically presented (Fig. 1B). We then manipulated the context around the occluded interval to investigate what information is required for restoration to occur.

**Figure 1.**
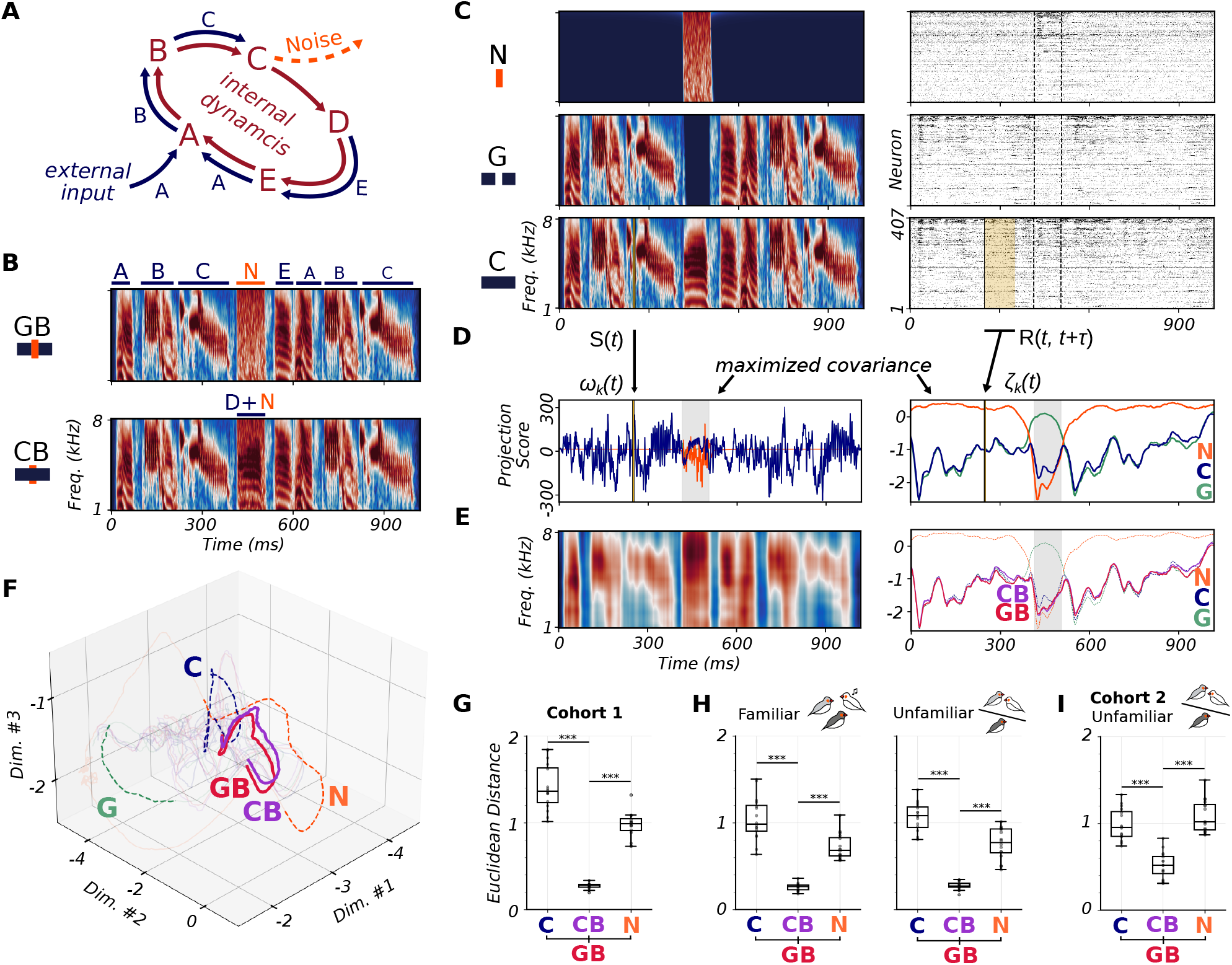
Neural responses to occluded stimuli. (**A**) Conceptual model: external input and internal network dynamics work together to drive the population through a sequence of states along a manifold, but when a syllable is occluded by noise the internal dynamics bias activity towards the state corresponding to the missing syllable. (**B**) Spectrograms of example illusion-inducing stimulus (top, Gap+Burst) and illusory percept (bottom, Continuous+Burst). See Fig. S1 for all motifs and critical intervals. (**C**) Spectrograms (left) and first cohort’s neural population response (*n* = 14 birds, 407 units) (right) to Continuous (unmodified), Gap (deleted syllable), and Noise (burst only) variants of the same motif in **B**; response sorted by units’ total absolute decoder coefficients. (**D**) Highest-ranked partial least squares component for spectrograms (left) and delay-embedded response (right) from **C**. (**E**) First latent dimension projections of CB and GB responses (withheld from training) (right), the latter used to decode GB’s stimulus spectrogram (left) (**F**) Projection of responses to example motif’s variants into the top three latent dimensions, highlighted for duration of CI. (**G**) Average Euclidean distance for all Cis (points: 8 motifs x 2 CIs) between GB and CB, C, and N. GB is closer to CB than to C (GLMM: *t*_38_ = 18.51, *p <* 0.001) or N (*t*_38_ = 11.38, *p <* 0.001). (**H**) First cohort data split by stimulus familiarity and the decoder was fit separately to each subset. For both conditions, GB was more similar to CB than to C or N (*F*_2,83_ = 207.75). No significant effect of stimulus familiarity on restoration (*F*_2,83_ = 0.048, *p* = 0.95). (**I**) GB trajectories were closer on average to CB trajectories than to either C (*t*_38_ = 7.30, *p <* 0.001) or N (*t*_38_ = 9.03, *p <* 0.001) in a separate cohort unfamiliar with all stimuli.

### Illusory neural responses to occluded syllables

We recorded the responses of 407 single units in the auditory pallium (*n* = 14 birds) to modified variants of eight zebra finch song motifs (stereotyped sequences of vocal elements) (Fig. 1B, C, Fig. S1A). Within each motif, we designated two non-overlapping critical intervals (CI) of 50–100 ms that spanned either an entire syllable or part of a syllable. For each critical interval, we created an illusion-inducing variant (GB: gap + noise burst) by replacing the sound in the interval with a white noise burst of the same duration, and a variant corresponding to the expected illusory percept (CB: continuous + burst) by adding white noise without deleting the underlying song (Fig. 1B). The amplitude of the noise was matched to the average amplitude of the motif ^23^. Two other variants were generated for each critical interval: a Gap variant (G) where the critical interval was deleted but not occluded by noise and a Noise variant (N) where the noise burst was presented on its own (Fig. 1C). We also presented the unmodified continuous motif (C). For this experiment, familiarity was controlled by socially housing half of the subjects with four of the eight vocalizers whose song motifs were used and half of the subjects in a separate room with the other four vocalizers. We predicted that restoration would occur only for the motifs with which each subject was directly familiar ^16^,^17^.

The effects of deleting syllables and replacing them with noise varied across the population of recorded units (Fig. 1C; Fig. S2). Instead of testing for restoration in individual neurons, we examined the responses of the whole population as trajectories within an eleven-dimensional latent subspace. We identified this subspace using partial least squares regression (PLS), a supervised method for dimensional reduction that combines aspects of principal components analysis and linear regression to find patterns of neural activity that maximally covary with the stimulus (Fig. 1C, D, Fig. S3A–C, see Methods). We structured this regression as a decoding model ^9^,^24^, with the stimulus at each moment in time predicted by the neural responses delay-embedded up to 100 ms into the future (because the delay between stimulus and response varied across the population). To avoid overfitting, the PLS model was trained using the C, G, and N variants for seven of the eight motifs and then tested on variants of the held-out motif, repeating this for all eight motifs.

Outside of the critical intervals, where the C, G, CB, and GB variants were the same, the neural trajectories were nearly identical (Fig. 1D, E). During the critical intervals (and immediately before, because of the delay-embedding), the trajectories for C, G, and N diverged. These observations confirm that PLS successfully identified latent dimensions of the population response that covaried with the stimulus and that different physical stimuli (syllable, gap, and noise) drove activity to different regions of the latent neural state space (Fig. 1D).

The key comparison for auditory restoration is between GB and CB during the critical interval. Despite their physical dissimilarity, these stimuli elicited nearly identical responses (Fig. 1E, Fig. S2C). In the first latent dimension, which has the highest covariance with the stimulus, the CB and GB trajectories often coincided midway between the trajectories for C and N, consistent with CBas a mixture of song and noise (Fig. 1E). The GB and CB trajectories remained close throughout the full eleven-dimensional latent subspace (Fig. 1F). To quantify this, we used the average Euclidean distance over time as a measure of similarity between the neural trajectories. The distances between GB and CB were consistently small for all the stimuli and critical interval locations, and consistently less than the distances to the corresponding C and N variants (Fig. 1G) regardless of interval location within the motif (Fig. S4C, D). In other words, even under anesthesia the auditory pallium responded the same whether the syllable in the critical interval was present or missing, as if it was responding to (or creating) the perceptual illusion ^10^.

Contrary to our expectations, there was no effect of familiarity, defined as having heard a specific bird’s song. The CB and GB trajectories were just as close in familiar and unfamiliar motifs, and just as far from the C and N trajectories (Fig. 1H). Because of this surprising result, we repeated the experiment in a second cohort of adult zebra finches (*n* = 10 birds, 398 units) who were not directly familiar with any of the eight song motifs, having been born after all of the vocalizers had died. Although the distances between trajectories are not directly comparable between cohorts because the number of units and their locations within the pallium were not the same, we observed the same pattern, with the GB trajectories much closer to CB than to C or N (Fig. 1I, Fig. S4E), confirming that restoration can occur even in the context of a song the bird has never heard before. We return to this point after a further test of the main hypothesis.

### Masking noise prevents context-based auditory predictions

Auditory restoration depends on context ^7^,^9^,^12^; thus, the similarity between replaced (GB) and added (CB) trajectories should depend on how clear the preceding syllables are. We tested this in a third cohort of birds (*n* = 5 birds, 398 units) by presenting two additional variants of the original eight motifs: one with white noise masking the entire motif (Continuous Masking, CM) and one with noise masking the motif with a deleted critical interval (Gap Masking, GM) (Fig. 2A). The masking noise was the same amplitude as the noise bursts. These two stimuli were physically identical to their respective counterparts, CB and GB, during the critical intervals, but we predicted the masking noise prior to the critical interval would deflect the neural trajectory away from the manifold, preventing the intrinsic network dynamics from restoring the missing syllables (Fig. 2B). As expected, masking the motifs pushed the neural trajectories away from the putative manifold defined by the response to C during the period leading up to the critical interval (Fig. 2C). Then, during the critical interval, the trajectories for the masked stimuli (CM and GM remained further away from C than the trajectories for the corresponding noise-burst stimuli (CB and GB) (Fig. 2C–E). In other words, the physically identical critical intervals (CB and CM; GB and GM elicited different responses depending on context: more like C when the context was clear and less when it was masked by noise.

**Figure 2.**
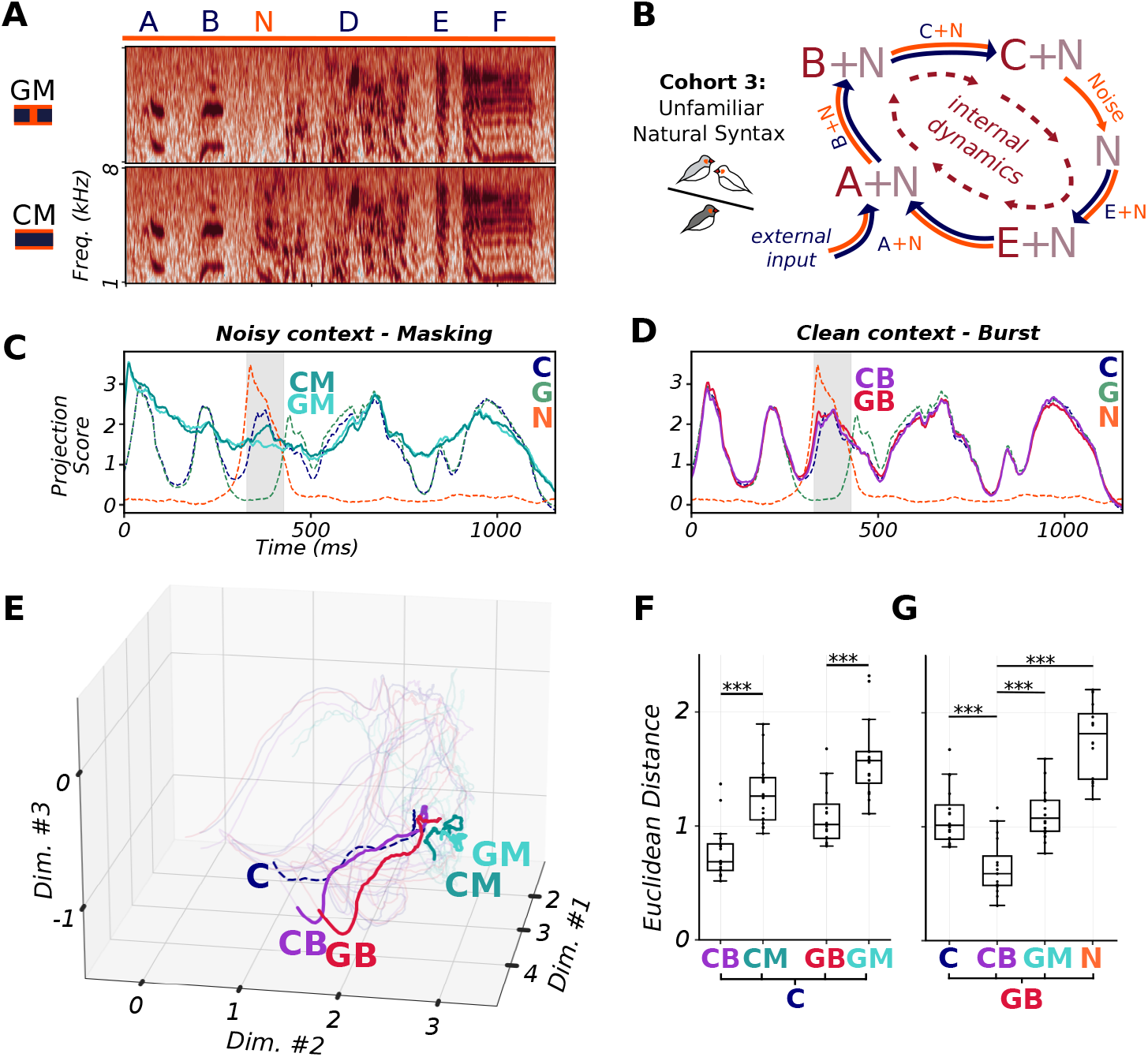
Neural responses to occluded stimuli versus noise-masked stimuli. (**A**) Spectrogram of example noise-masked stimuli variants: gap + noise masking motif (GM, top) and continuous + noise masking motif (CM, bottom). (**B**) Conceptual model: masking noise drives the population off the learned manifold to regions of state space where the dynamics do not restore the missing syllable. (**C**) First latent dimension of CM and GM responses, plotted against same-motif trajectories C, G, and N. GM and CM trajectory frequently diverge from C trajectory both during and outside of CI. (**D**) CB and GB trajectories were nearly identical to the song trajectory C, even during critical interval. (**E**) Projection of responses to noise-burst (CB and GB) and noise-masked (CM and GM) variants alongside the putative song trajectory C into the top three latent dimensions, highlighted for duration of CI. (**F**) Average Euclidean distances to unmodified song C across CIs between pairs of burst-masking variants. Burst variants CB and GB are closer to song trajectory C than masking variants CM (paired t-test: *t*_15_ = 9.82, *p <* 0.001) and GM (*t*_15_ = 9.83, *p <* 0.001), respectively. (**g**) GB responses closest to illusory percept trajectory CB compared to control trajectories C (GLMM: *t*_53_ = 4.77), GM (*t*_53_ = 5.01), or N (*t*_53_ = 11.96).

As in cohorts 1 and 2, the GB trajectory was closer to CB during the critical interval than to the trajectories for any of the other stimuli, including GM (Fig. 2F, Fig. S4F). Like N, GM is physically identical to GB during the critical interval and can therefore serve as a reference for the trajectory we might expect if restoration were not occurring. Unlike N, GM does not have abrupt acoustical onsets or offsets, which produced transient responses (Fig. 1D, 2C) that contributed to the dissimilarity between GB and N trajectories. GM is an important control, because CB and GB trajectories might look like each other for two reasons: either the context of GB induces a response to the missing syllable (restoration), or the noise in CB overwhelms the response to the underlying syllable (masking). Thus, the observation that CB and GB trajectories were similar to each other but not to N or GM is consistent with restoration, not masking.

### Natural song syntax is required for restoration

How can the auditory pallium fill in parts of songs the bird has never heard before? According to our model, repeatedly hearing the sequence of elements in a song entrains local dynamics (Fig. 1A), but we observed restoration for unfamiliar songs in three separate experiments (Fig. 1H–I, 2F). One possibility is that experience shapes dynamics to encode not just the specific sequence of elements in specific songs, but more general transition probabilities between syllable types that might be shared among birds in our colony or more broadly within the species. In support of this idea, zebra finch song elements from geographically separate populations cluster into about 10 loosely defined categories, and there are consistent biases in how element types are ordered within songs ^25^,^26^. These statistical regularities could shape intrinsic dynamics through experience, biasing local circuitry towards regions of the neural state space corresponding to plausible candidates for successive syllables.

If this hypothesis is correct, then we expect that restoration will not occur for stimuli that do not follow the typical syntactic structure of zebra finch song (Fig. 3A, B). To test this prediction, we created a set of eight synthetic motifs, each a sequence of five unique syllables chosen at random from 60 syllables extracted from another colony’s song recordings (Fig. 3A, Fig. S1B). We did not alter any of the individual syllables but randomized their order in the motif, disrupting the natural transition probabilities that we hypothesize are necessary for context-based syllable prediction (Fig. 3B). As with the natural motifs, we designated two critical intervals in each scrambled motif, one in the second syllable and one in the fourth. The critical intervals were all 70 ms in duration, and we placed them 10–40 ms after the start of the syllable. This was to provide a stronger test of the role of syntax, because the syllable identity could potentially be predicted from the initial non-occluded segment without the context of conspecific syntax. For each motif and critical interval, we constructed C, G, N, CB, GB, CM, and GM variants as before. We collected single unit responses to these stimuli in separate recordings from the same third cohort of birds (*n* = 5 animals) and used PLS to identify a separate 11-dimensional latent subspace for these stimuli and responses.

**Figure 3.**
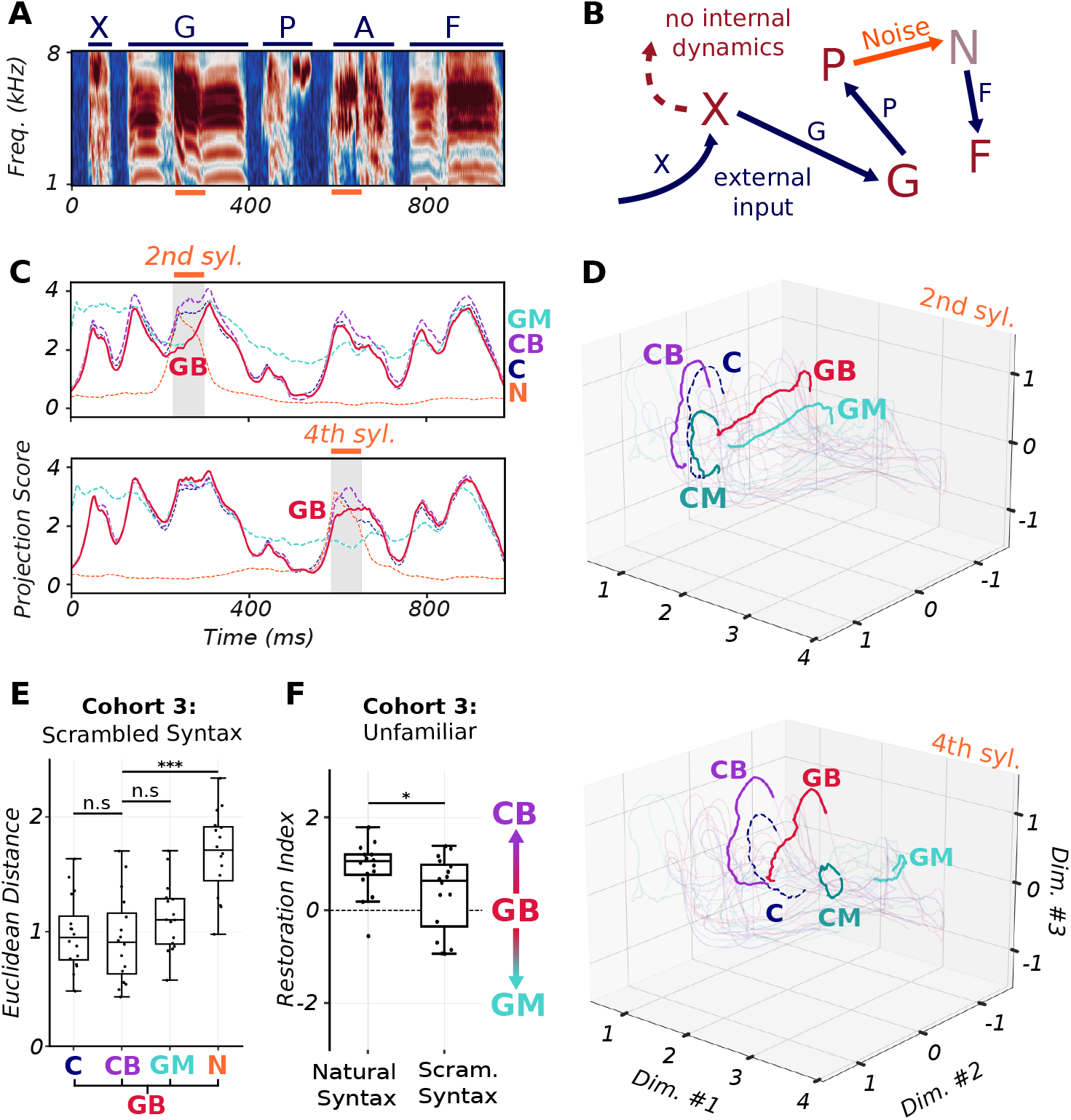
Neural responses to scrambled syntax stimuli. (**A**) Spectrogram of example scrambled syntax motif (all motifs in Fig. S1). Critical intervals indicated by horizontal orange bars. (**b**) Conceptual model: no internal dynamics corresponding to transitions in scrambled motifs, response determined only by stimulus features. (**c**) First latent dimension of GB (solid line) plotted against trajectories for C, CB, GM and N variants (dashed lines) for second (top) or fourth (bottom) syllable occlusion. GB trajectory coincides with GM during first interval (top), half-way between CB and GM in second interval (bottom). (**d**) Top three latent dimensions of neural response as in (C). GB resembles non-syllable trajectory GM compared to syllable presentations C, CB, and CM for second syllable occlusion (top) while resembling CB trajectory in fourth syllable occlusion (bottom). (**e**) Average Euclidean distance from GB to control variants C, CB, GM, and N for all scrambled syntax critical intervals. GB—CB distances not significantly different from GB-–GM (GLMM: *t*_53_ = 1.55, *p* = 0.127) or GB–C (*t*_53_ = 0.40, *p* = 0.694), but smaller than GB—N distances (*t*_53_ = 6.20, *p <* 0.001). (**f**) Restoration index (see Methods, Fig. S3E) scores for natural and scrambled syntax motif from cohort 3 neural responses indicate if GB trajectory closer to CB(positive RI) or GM (negative RI). Natural motif RIs significantly higher than zero (one sample t-test: *t*_15_ = 5.81, *p <* 0.001) and higher than scrambled syntax RIs (paired t-test: *t*_15_ = 2.78, *p* = 0.014).

In contrast to natural motifs, the trajectories for GB variants of scrambled motifs were dissimilar to their CB counterparts. Indeed, for some critical intervals, the GB trajectory was closest to GM, which is physically identical to GB during the critical interval (Fig. 3C, D). On average for all critical intervals of scrambled motifs, GB trajectories were no closer to CB than to GM (Fig. 3E, Fig. S4H), indicating that restoration was weaker or absent without natural song syntax. To compare the restoration effect between natural and scrambled syntax motifs within the cohort, we calculated a restoration index (RI; Methods, Fig. S3E) based on the average distances between GB, CB, and GM in a critical interval. For natural (unfamiliar) motifs, RI was greater than zero and greater than for scrambled motifs (Fig. 3F).

### Illusory responses are distributed throughout the auditory pallium

Recordings in the third cohort of birds used higher-density probes with more channels, resulting in high enough yields to test for restoration in individual birds and recordings. Subdividing the data by animal produced the same result: RI for natural stimuli was both greater than zero and greater than RI for scrambled motifs (Fig. 4A, B).

**Figure 4.**
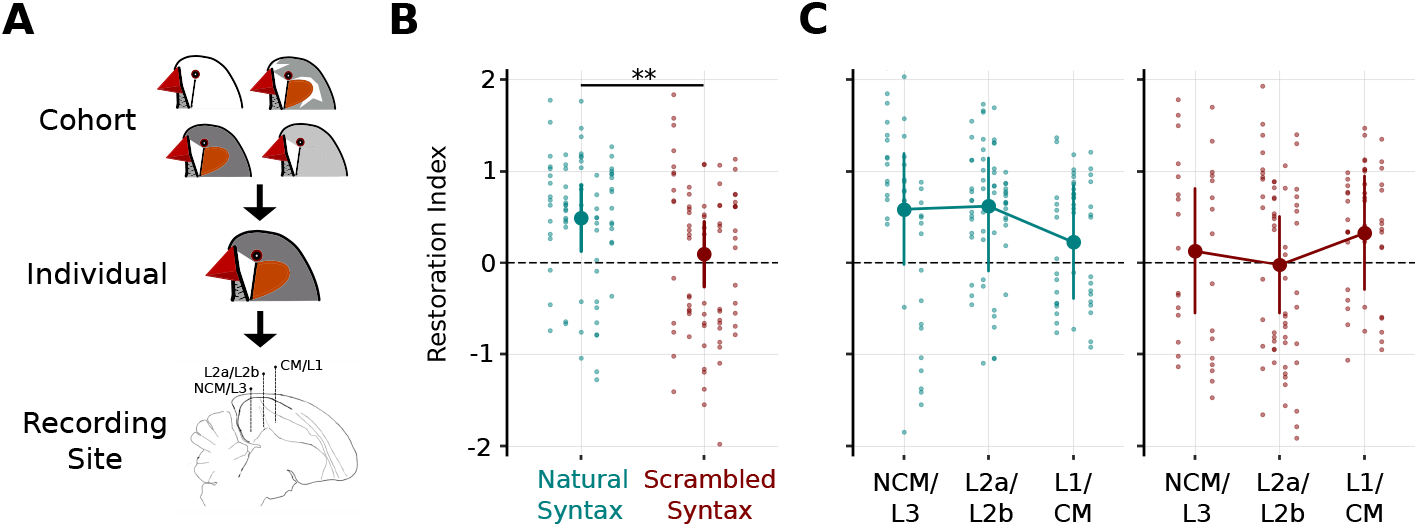
Auditory restoration for individuals and pallial regions. (**A**) Hierarchy of neural data to which the partial least squares model was applied: from the cohort-pooled neuron population to individual subject’s neurons, to acute recording sites grouped by auditory pallium region. (**b**) Restoration Index (RI) scores for natural motifs versus scrambled motifs for individual birds in cohort 3. Each dot represents a critical interval’s RI. Each column of dots corresponds to a subject. Filled circles show estimated mean RI for stimulus set; whiskers show 95% confidence intervals for the mean. RI scores are lower for scrambled syntax than for natural syntax (GLMM: *t*_154_ = 3.3, *p <* 0.01), and RI is only positive for natural songs (*t*_5.8_ = 3.4; *p* = 0.016). (**c**) RI scores for natural motifs (left) and scrambled motifs (right) in each recording site: NCM/L3 (*n* = 3 recordings), L2 (*n* = 4 recordings), and L1/CM (*n* = 3 recordings). As in (B), but with columns corresponding to individual sites. There was no significant main effect for region (*F*_2,11.85_ = 0.036, *p* = 0.96) or stimulus set (*F*_2,11.09_ = 3.2, *p* = 0.10), or for the interaction (*F*_2,11_ = 1.6; *p* = 0.25), but the average of RI across areas was greater than zero (*t*_5.8_ = 3.4; *p* = 0.016).

We subdivided the data further by recording site to determine if the illusory activity originated in a specific area. The avian auditory cortex comprises several laminar subdivisions organized in a hierarchy of increasingly complex functional properties ^27–29^ and it is possible that the illusion emerges in a similar hierarchical fashion. To test this hypothesis, we grouped our recordings into three areas, the thalamorecipient areas L2a and L2b (*n* = 4 sites for 3 birds), the superficial areas CM and L1 (*n* = 3 sites for 2 birds), and the deep/secondary areas NCM and L3 (*n* = 3 sites for 2 birds; note that recordings were obtained from multiple areas in some animals). Contrary to our prediction, RI did not differ significantly among regions for either natural or scrambled stimuli (Fig. 4C), indicating that illusory activity was present throughout the pallium.

## Discussion

Perceptual illusions present a unique opportunity to investigate how the brain uses internal models in sensory processing. Normally, internal models are aligned with incoming sensory input; but in an illusion, perception diverges from reality, and the neural activity produced by a model can be dissociated from the activity produced by the physical stimulus. In this study, we observed that the zebra finch auditory pallium responds to noise-occluded songs as though the missing syllables are present, even when birds are under anesthesia and have never heard the song before (Fig. 1). These results suggest that local dynamics are sufficient to bias auditory responses towards regions of the neural state space corresponding to missing parts of the song. This bias is weaker if the song is masked by noise (Fig. 2) or if the order of the preceding syllables is disrupted (Fig. 3), indicating that the dynamics reflect general acoustical and syntactic features of song rather than specific memories.

Applying dimensional reduction methods like partial least squares to population activity in sensory systems reveals trajectories along manifolds in the neural state space that depend on the features of the stimulus ^30^,^31^ and other task variables ^32^,^33^, but it can be difficult to determine to what extent the trajectories reflect ascending sensory drive versus intrinsic network dynamics. Our results show that the trajectories for GB and CB remain closely aligned despite the physical stimuli being different, thereby uncovering intrinsic circuit dynamics underlying the manifold. Importantly, the trajectories only stay aligned for unmasked songs with natural syntax, indicating that local dynamics only override sensory input when the population is on the manifold. Taken together, the results are suggestive of a dynamical system in which the vector field is weak and disorganized except near the manifold, where it creates a flow towards and along a defined trajectory.

The zebra finch auditory pallium exhibits a hierarchy of selectivity for increasingly complex features of song ^27^, and individual neurons in higher-order areas exhibit sensitivity to context ^22^,^34^ consistent with our hypothesized dynamics. We observed illusory activity not just in the higher-order areas but throughout the pallium (Fig. 4), suggesting that it emerges from the dynamics of the whole recurrently connected network ^18^,^20^. By strengthening connections between ensembles of neurons sequentially activated by conspecific songs, Hebbian plasticity ^35^ is well-suited to create these dynamics, but to our surprise, it was not necessary for birds to have heard a specific sequence of syllables for the pallium to fill in missing segments. We interpret this result to mean, firstly, that the pallium responds not only to the low-level acoustical features of syllables but their categories as well. There are only around 10 different types of syllables in zebra finch song ^25^,^26^, and auditory responses in higher-order pallial areas show evidence of category selectivity ^28^,^36^. Secondly, it implies that the dynamics do not encode deterministic sequences but rather the transition probabilities between syllable categories, which are not uniform ^26^. This could be instantiated as branch points on the manifold or as a superposition of multiple candidates, as proposed for human speech perception by the Trace model and its successors ^4^,37. In these models, top-down modulation would further bias the dynamics towards the most contextually appropriate candidates ^38^,^39^, effects that should become more apparent on a trial-by-trial basis when birds are awake and engaged in a task.

Genetic factors may also contribute to an internal model for the general syntactic structure of zebra finch song. Like other songbirds, male zebra finches require a tutor to produce normal song but have innate preferences for species-typical syllables and sequences ^40^,^41^ that are at least partially perceptual ^42^. Investigating restoration in finches raised in acoustically impoverished environments ^43^, may begin to elucidate the neural basis of these poorly understood genetic preferences and how they interact with experience. More broadly, explicitly modeling the dynamics of sensory systems during perceptual illusions has the potential for new insights into the mechanisms of the brain’s internal models and their role in decoding noisy sensory inputs.

## Materials and Methods

### Animals

All animal use was performed in accordance with the Institutional Animal Care and Use Committee of the University of Virginia. Adult zebra finches were obtained from the University of Virginia breeding colony. Eight zebra finches (all male) were used to provide song stimuli and for song familiarization for the natural song stimuli set. For the first cohort, 8 zebra finches (all male) were used for song familiarization, and 14 (7 female) were used for extracellular recording. For the second and third cohorts, 10 (6 females) and 5 (3 females) were used for extracellular recording, respectively.

### Song stimuli and social familiarization

Acoustic recordings of song were made by housing each singer individually in a sound isolation box (Eckel Industries, Cambridge, MA) with *ad libitum* food and water on a 16:8 h light:dark schedule. A lavalier microphone (Audio-Technica Pro 70) was positioned in the box near a mirror to stimulate singing. The microphone signal was amplified and digitized with a Focusrite Scarlett 2i2 at 44.1 kHz, and recordings to disk using custom C++ software (jill: https://github.com/melizalab/jill; version 2.1.4) were triggered every time the bird vocalized. A typical recording session lasted 1–3 days. From each bird’s recorded corpus, a single representative song motif of 837–1200 ms (mean±SD: 998.9 ± 130.6 ms) was selected, high-pass filtered using a 4th-order Butterworth filter with a cutoff frequency of 500 Hz.

One cohort of 14 birds (7 female) was socially familiarized with half of the songs in the stimulus set. The eight singers were semi-randomly assigned to two groups of four after we recorded their songs, taking care that songs were dissimilar between groups (Fig. S1A). Each group was housed together in separate, acoustically isolated rooms within the breeding colony. Each room housed 20–40 other adult and juvenile zebra finches. For at least one week before electrophysiology, experimental birds were housed in the same cage with one of these groups. Subjects were never housed in the same or adjoining cages with the unfamiliar singers. Thus, familiarity was counterbalanced: half of the subjects were closely familiar with one half of the songs, and the other half of the subjects were familiar with the other half of the songs.

For each of the eight motifs, we designated two non-overlapping critical intervals to test for restoration. Critical intervals began at the onset of a syllable at least 113 ms after the start of the motif. They were 58–100 ms in duration (mean ± SD: 83 ± 16 ms) and ended at least 263 ms before the end of the motif. For each motif and critical interval, we generated four variants (Fig. 1B, C). In the gap variant (G), the critical interval was deleted and replaced with silence. In the gap-burst variant (GB), the critical interval was deleted and replaced with white noise equal to the average amplitude of the motif. In the continuous-burst variant (CB), the same white noise burst was added to the critical interval without deleting the underlying song. The noise variant (N) consisted of the noise burst by itself. We also presented the unmodified continuous motif (C), which was the same for both critical intervals. A 3-ms cosine ramp was applied to the beginnings and ends of the motifs, the gaps, and the noise bursts to avoid discontinuities in the digital signals.

Birds in the second and third electrophysiology cohorts were born after all of the singers had died. Thus, it was impossible for them to have been directly familiar with any of the songs, but they may have been familiar with songs from birds who had directly or indirectly copied some of the elements or sequences from the original singers. We do not record the songs of every bird in our colony and were unable to confirm whether this was the case. Two additional variants were presented to birds in these cohorts, both created by adding white noise spanning the entire motif. In the gap-masked variant (GB), the critical interval was deleted prior to adding masking noise. In the continuous-masked variant (CM), the masking noise was added to the unmodified motif. The masking noise was the same amplitude as the noise burst, and a 25-ms cosine ramp was applied to the beginning and end.

The third cohort was also presented with stimuli derived from “pseudo-motifs” comprising synthetic sequences of zebra finch syllables. A total of 68 different syllables were extracted from acoustic recordings of 16 birds made in the Margoliash lab (University of Chicago) that were filtered and normalized the same way as the recordings from our lab. Eight unique, non-overlapping sequences of 5 syllables were selected without replacement from this pool, with silent gaps of 30 ms separating the syllables. Two critical intervals were designated for each pseudo-motif in the second and fourth syllables, therefore these two syllables from each motif were selected from the reduced pool of syllables that were longer than 100 ms. Syllable selection was pseudo-randomly selected (seeded) using the `numpy` python library. The critical intervals 70 ms in duration were placed at least 10 ms after the start of the syllable. C, G, N, GB, CB, GM, and CM variants were generated for each critical interval as described above. For these scrambled syntax stimuli, a 2-ms cosine ramp was applied to the beginnings and ends of the syllables, the gaps, and the noise bursts.

All of the stimulus manipulations were performed using custom Python code (TBD; to be released after initial review). Single-precision floating point arrays were used throughout the manipulations to preserve dynamic range, and all of the motifs within each stimulus set were normalized to the same RMS amplitude (–27 dB relative to full scale for cohorts 1 and 2; –30 dB for cohort 3) prior to processing to prevent clipping.

### Extracellular recordings

#### Surgery

Birds were anesthetized with isoflurane inhalation (1–3% in O2) and placed in a stereotaxic apparatus (Kopf Instruments). An incision was made in the scalp, and the skin was retracted from the skull. The recording site was identified using stereotaxic coordinates relative to the Y-sinus. A metal pin was affixed to the skull rostral to the recording site with dental cement, and the skull over the recording site was shaved down but not completely removed. The bird was allowed to recover completely for several days prior to recording.

On the day of recording, the bird was anesthetized with three intramuscular injections of 20% urethane spaced half an hour apart. The bird was placed in a 50 mL conical tube, and the head pin was attached to a stand in the recording chamber. The thin layer of skull remaining over the recording site was removed along with the dura, and a well was formed around the recording site and filled with phosphate-buffered saline.

#### Stimulus presentation

Stimuli were presented with the `sounddevice` python library (version 0.3.10) through a Samson Servo 120a amplifier to a Behringer Monitor Speaker 1C. The amplifier gain was adjusted to give an RMS amplitude for the unmodified motifs of 70 dBA SPL at the position of the bird’s head. Stimuli were presented in a pseudorandom order to minimize stimulus adaptation, with 1 s between each song. Each stimulus was presented 10 times.

#### Data acquisition

In cohort 1, neural recordings were made using a NeuroNexus 32-channel probe in a four-shank, linear configuration (A4×8-5mm-100-400-177-A32) connected to an Intan RHD2132 Amplifier Chip. In cohorts 2 and 3, recordings were made using a two-shank 128-channel probe (128 P and 128 DN, Masmanidis Lab) connected to an Intan RHD2164 Amplifier Chip, or a two-shank 64-channel probe (HSSY-H6, Cambridge Neurotech) connected to two Intan RHD2132 Amplifier Chips. For all recordings, data were collected from the Intan chips using an Open Ephys Acquisition Board and sent to a computer running Open Ephys GUI software (version 0.4.6).

The recording electrode was coated with DiI (Invitrogen). In cohort 1, the electrode was inserted at a dorso-rostral to ventro-caudal angle that allowed for recording of all auditory forebrain regions with a single penetration. In cohorts 2 and 3, the probe was inserted at a more vertical angle to confine the recording sites to superficial areas (L1 and CM, the caudal mesopallium), the intermediate thalamorecipient areas (L2a and L2b), or the deep/secondary areas (L3 and NCM, the caudo-medial nidopallium) The probe was lowered into the brain until the local field potentials across channels and shanks showed coordinated responses to birdsong, and the probe was allowed to rest in place for half an hour to avoid drift during the recording. After each recording, the probe was advanced far enough so that none of the recording sites overlapped with the previous position of the probe, and additional recordings were made until auditory responses were no longer observed, at which point we withdrew the probe and either moved to a new location or terminated the experiment. In subjects presented with both stimuli sets (natural vs. synthetic syntax), a recording of each set was performed separately at each recording site. The order of the stimulus sets was alternated at successive sites.

#### Histology

After recording, birds were administered a lethal intramuscular injection of Euthasol and perfused transcardially with a 10 U/mL solution of sodium heparin in PBS (in mM: 10 Na2HPO4, 154 NaCl, pH 7.4) followed by 4% formaldehyde (in PBS). Brains were immediately removed from the skull, postfixed overnight in 4% formaldehyde at 4 °C, cryoprotected in 30% sucrose (in 100 mM Na2HPO4, pH 7.4), blocked saggitally into hemispheres or on a modified coronal plane ^44^, embedded in OCT, and stored at –80 °C. 60 µm sections were cut on a cryostat and mounted on slides. After drying overnight, the sections were rehydrated in PBS and cover-slipped with Prolong Gold with DAPI (ThermoFisher, catalog P36934; RRID:SCR_015961). Sections were imaged using epifluorescence with DAPI and Texas Red filter cubes to located DiI-labeled penetrations. Images of the electrode tracks were used to identify the locations of recorded units based on where the recording site associated with each channel was located. In cohort 1, units were located in the caudal mesopallium (CM, *n* = 56 units); field L subunits L1 (*n* = 25 units), L2a (*n* = 33 units), and L3 (*n* = 59 units); and the caudomedial nidopallium (NCM, *n* = 90 units). In cohort 3, because electrodes were inserted dorsal-ventrally, we assigned each identified recording site to one of three broader groups of areas: L1 and CM (*n* = 3 recordings), L2a and L2b (*n* = 4 recordings), NCM and L3 (*n* = 3 recordings).

#### Spike sorting

Spikes were sorted offline. Recordings from cohort 1 were sorted using MountainSort 4; recordings from cohorts 2 and 3 were sorted with Kilosort 2.5. We used `phy` to further curate single units by visual inspection for spheroid PCA cluster shape, very low refractory period violations in the autocorrelogram, and stability of the unit throughout the recording. For cohort 1, single units were only included in the dataset if they showed a clear, phase-locked auditory response to at least one stimulus. For cohorts 2 and 3, we used all well-isolated units. In cohort 3, the recordings for natural and synthetic stimulus sets at the same site were sorted separately.

#### Neural manifold decoding

Our goal in this analysis was to identify a low-dimensional latent subspace in which to compare the responses of the neural population to the illusion-producing stimulus (GB) and to the presumed illusory stimulus (CB) as well as other variants. A dimensionality reduction technique was needed to determine the relevant auditory components of neural activity, due to the diverse neuronal firings found across critical intervals and song motifs (Fig. S2). Principal components analysis (PCA) is often used to find dimensions of neural activity that explain the highest variances in the population response, but these dimensions are not necessarily informative about the stimulus. Therefore, we turned to partial least squares (PLS) regression, which simultaneously finds dimensions in the stimulus and response that are maximally covariant. We formulated this regression as a decoding model, in which each frame in the stimulus spectrogram **Y**_*t*_ is a (linear) function of the population response over a window from *t* to *t* + *τ*.

#### Stimulus spectrogram

The sound pressure waveforms of the stimuli were converted to time-frequency representations using a gammatone filter bank, implemented in the Python package `gammatone` (version 1.0) with 50 log-spaced frequency bands from 1–8 kHz, a window size of 2.5 ms, and a step size of 1 ms. Power was log-transformed with a constant offset of 1, giving the transformed signal a lower bound of 0 dB. This produces the *n*_*t*_ *× n*_*targets*_ stimulus matrix **Y**, where *n*_*targets*_ is the number of frequency bands and *n*_*t*_ is the number of spectrogram frames.

#### Response matrix

The responses of each unit *i* were averaged across all 10 trials and binned at 1 ms resolution. The resulting response vector was delay-embedded 100 ms into the future to give an *n*_*t*_ *×* 100 Hankel matrix **R**_*i*_, and then projected into a basis set comprising 15 raised cosines (^45^) to get an *n*_*t*_ *× n*_*basis*_ matrix **X**_*i*_. The basis vectors were spaced log-linearly and increased in width with lag, which gives the model high temporal resolution at short lags and lower resolution at longer lags. This allows the inclusion of longer lags without exploding the number of parameters. All of the units within each analysis (initially each cohort, but later by subject and recording) were pooled by concatenating **X**_*i*_ along the columns, resulting in an *n*_*t*_ *× n*_*f eatures*_ response matrix **X**, where *n*_*f eatures*_ = *n*_*units*_*n*_*basis*_ and *n*_*t*_ is the number of time points (Fig. S3A).

#### Partial least squares regression

To estimate the linear coefficients of the decoder model, we used the supervised cross composition method Partial Least Square implemented as `PLSRegression` in the python package `scikit-learn` (version 1.3.2) (^46^). As PLS is not commonly used in systems neuroscience, we provide a brief explanation of the procedure here. Through an iterative process, the PLS algorithm finds *k* pairs of orthogonal weight vectors *µ*(*t*) and *ν*(*t*) for **X** and **Y** respectively, such that *k ≤ min*(*n*_*f eatures*_, *n*_*targets*_) and the respective projections of **X** and **Y** – the score vectors *ξ*(*t*) and *ω*(*t*) – are maximized in covariance. At each iteration, weight vectors are calculated as the first left and right singular vectors of the covariance matrix **X**^*⊤*^ **Y**, followed by the score vectors by projecting **X** and **Y** onto the weights. A loading vector *γ*(*t*) is found such that the product *ξ*(*t*) *· γ*(*t*)^*⊤*^ approximates **X**. This product is subsequently subtracted from **X** for the next iteration. Of note, the predictive model of PLS regression has the loading vectors *δ*(*t*) for **Y** calculated using the response matrix **X**’s score vector rather than **Y**’s score vector. This step ultimately affects the low-dimensional projection of **Y** such that the percentage of explained variance cannot be calculated for out-of-sample stimulus spectrogram matrices as can be done for response data (Fig. S3B). A projection (or rotation) matrix **P** is calculated from the weight and loading matrices, **M** and **Γ**, such that the score matrix **Ξ** = **X** *·* **P** for training data. **P** allows us to project out-of-sample neural responses 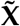 into trajectories 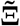 within the latent subspace (Fig. S3C, D). One can also calculate a coefficient matrix *β* to predict the stimulus from the response 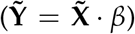 for the purposes of optimizing model parameters; we otherwise confine our analysis to the latent neural subspace.

#### Model parameters

Model coefficients were estimated with data from C, G, N, and CM (when available) stimulus variants, leaving out GB, CB, and GM (when available) for later testing. We used 8-fold cross-validation (holding out one motif per fold) to determine the best values for the number of components *k*, the number of time lags *τ*, the number of raised-cosine basis functions *q*, and a linearity parameter that controls the spacing of the basis functions. The *τ, q*, and linearity parameters were estimated from modeling the first cohort neural population, and held constant for all subsequent models and cohorts: *τ* was 100 bins at 1-ms resolution, the number of basis functions was 15, and the linearity factor was 20. Cross-validation was thus used to find the optimal *k* number of latent dimensions for each neuron population:

Neural data from the first cohort (407 units) were first analyzed together (Fig. 1G), and subsequently separated by familiarity (*n* = 191 units and *n* = 216 units, Fig. 1H) for modeling. All three models reached optimal decoding performance at *k* = 11 components, which explained 11.085 ± 0.95% (full cohort), 13.93 ± 1.27% and 12.62 ± 0.73% (split cohort) of training data variance. The second cohort’s model reached optimal performance at *k* = 12 components, with 10.96 ± 0.58% variance. However, we constrained the analysis of the second cohort to 11 components for comparison with first cohort’s result (Fig. 1I). For the third cohort, the separately-trained models for natural and scrambled syntax stimuli reached optimal performance at *k* = 11 components, explaining 14.59 ± 0.34% and 14.65 ± 0.42% of training data variance respectively. For determining the effects of restoration across the zebra finch pallium, neural data was first partitioned by individual subjects for separate PLS modeling (Fig. 4B), then by individual acute recording sites (Fig. 4C). We do not report each trained model’s best fit parameter nor performance here, as they vary widely between recordings.

#### Analysis of neural trajectory

After choosing hyperparameters, we analyzed each motif by fitting the PLS model to C, G, N, and CM from the other 7 motifs and then transforming the response to each variant of the eighth motif 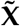 into a latent trajectory 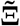 within a *k*-dimensional subspace. We calculate the similarity of two responses as the Euclidean distance between two trajectories at each point in time, averaged over time. For example, the similarity between responses to GB and CB is

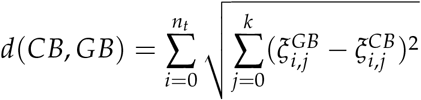

Four distances are relevant to assessing auditory restoration, between the illusory stimulus GB and the control variants C, CB, N, and GM (when available). A generalized linear mixed-effect model (GLMM) was used to infer each control variant’s average distance to GB within each model instance:

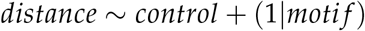

The variable *distance* depended on the *control* condition (coded as a factor with 4 levels for each control variant) with a random intercept for each *moti f* (coded as a factor with 8 levels). Each partial least squares modeling instance provided 16 observations, corresponding to 16 critical intervals in either stimulus set. In the case of cohort 1 where familiarity was internally controlled, the binary variable *f amiliarity* can be added as an interaction term to the GLMM: 1 if the motif from which the distance measures were taken was familiar to the model’s subjects, 0 if unfamiliar.

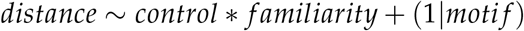

#### Restoration index

We calculated a restoration index to score the relationship between the GB trajectory and its controls, CB and GM. Within a critical interval, we used the Euclidean distance averaged over the entire length of the interval. The restoration index formula is as follows:

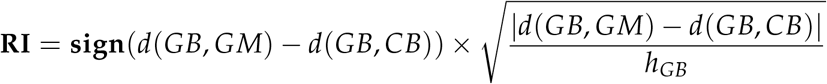

where the neural states GB, GM, and CB each were treated as a vertex of a triangle within the n-dimensional space of the projection. GB’s proximity to either CB and GB gives the RI arbitrary sign – here a positive RI means GB is closer to CB than to GM. Using the averaged distances as the side lengths (Fig. S3E), we scaled the difference by the altitude *h*_*GB*_ from vertex GB as a measure GB’s overall proximity to both CB and GM. *h*_*GB*_ can be calculated using Heron’s formula.

For the third cohort, where a single model was fitted to the pooled population response for natural versus scrambled motifs separately, comparison of RI scores between the two models were done with a paired t-test – here equivalent to a GLMM. Additional sources of variance introduced by modeling data from individual subjects or acute recording sites thus required fitting a mixed effects model. In the case of subject-level comparison between natural and scrambled motifs:

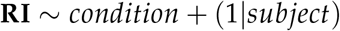

The binary variable *condition* was coded 0 for scores from the natural stimulus model, and 1 for scores from the scrambled syntax model. Similarly, the area-level analysis relied on the formula:

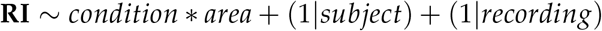

where the variable *area* had 3 coded factors for the combined auditory pallium regions NCM/L3, L2a/b, and CM/L1. Random effects from *subject* and *recording* accounted for different sources of variance.

Unless otherwise noted, all comparisons were fit using the `lmer` function from the R package `lmerTest` (version 3.1.3), which wraps around the homonymous function from `lme4` (version 1.1.35.3). Calculation of estimated marginal means and joint hypothesis tests were done using the package `emmeans` (version 1.10.4).

## Data and code availability

The electrophysiological data that support the findings of this study are available in figshare with the identifier TBD (to be supplied after initial review of manuscript). The custom R and Python code used in this study are available on github at TBD (to be supplied after initial review).

## Author Contributions

MCB, CDM, and BL conceived and planned the experiments. MCB and BL carried out the experiments. BL analyzed the data, and prepared the figures. BL, CDM, and MCB wrote the manuscript.

## Additional Information

Correspondence and requests for materials should be addressed to CDM.

## Supplemental Data

Supplementary Figures 1–4

**Supplemental Figure 1.**
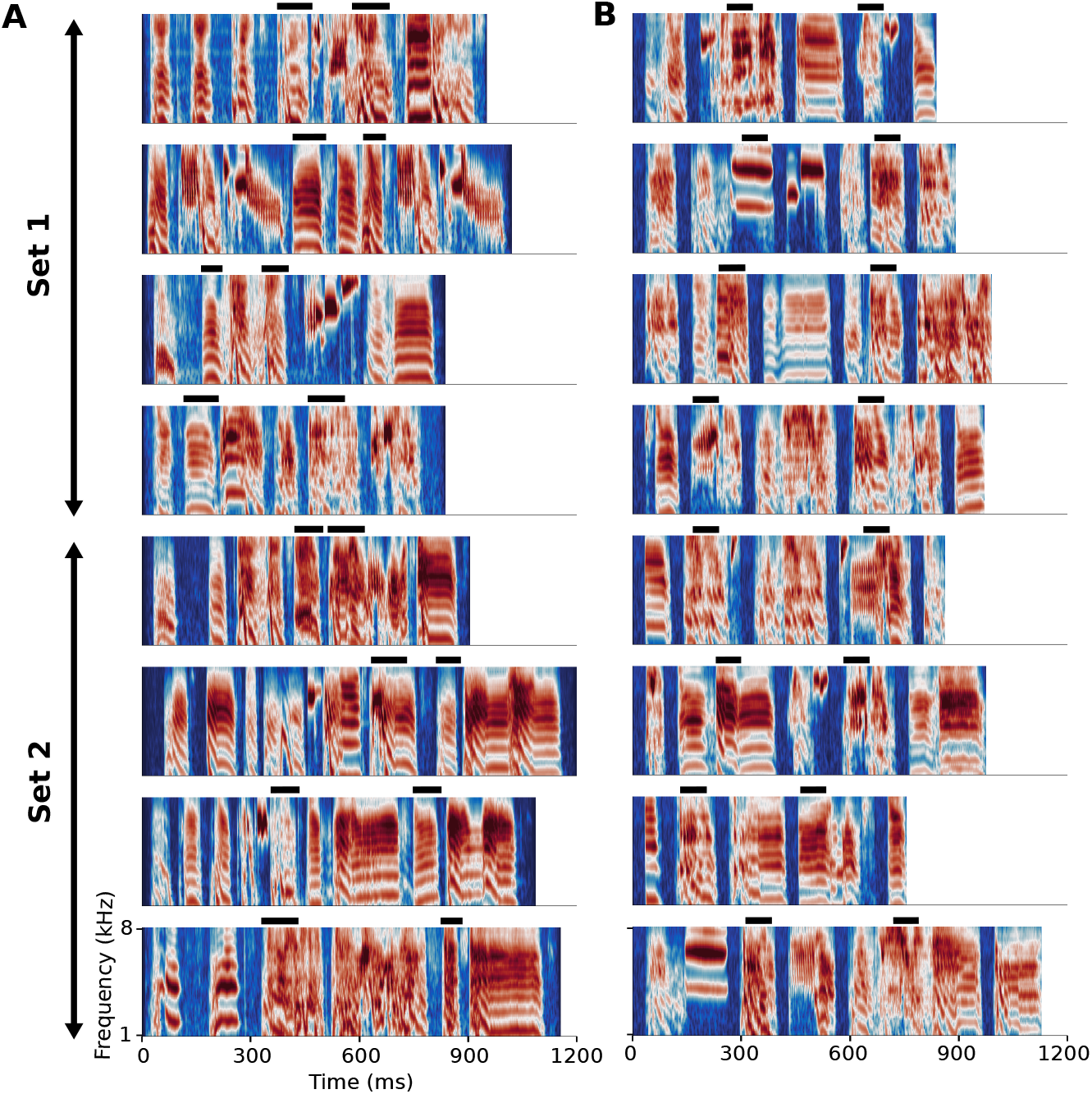
Song motifs used in electrophysiology recordings. (**A**) All eight song motifs used in the natural stimuli set. Each panel is a gammatone spectrogram, which has logarithmically spaced frequency bands spanning from 1–8 kHz. Stronger shades of red-orange indicate greater power. Each black bar above a panel indicates location of critical interval within the motif. Subjects from the first cohort were reared in the same colony as four of the eight vocalizers, thus familiar with song motifs from set 1 (top four song motifs) and unfamiliar with motifs from set 2 (bottom four song motifs), or vice versa. (**B**) Gammatone spectrograms of all eight synthetic song motifs used in the scrambled syntax stimuli set. Each scrambled motif contains five randomly-chosen syllables from a pool of 60, with the second and fourth syllables at least 100 ms long. Each syllable in the motif set was unique, although synthetic motifs may share similar syllables. Critical intervals of 70 ms were placed within either the second or fourth syllable, indicated by black bars above spectrograms.

**Supplemental Figure 2.**
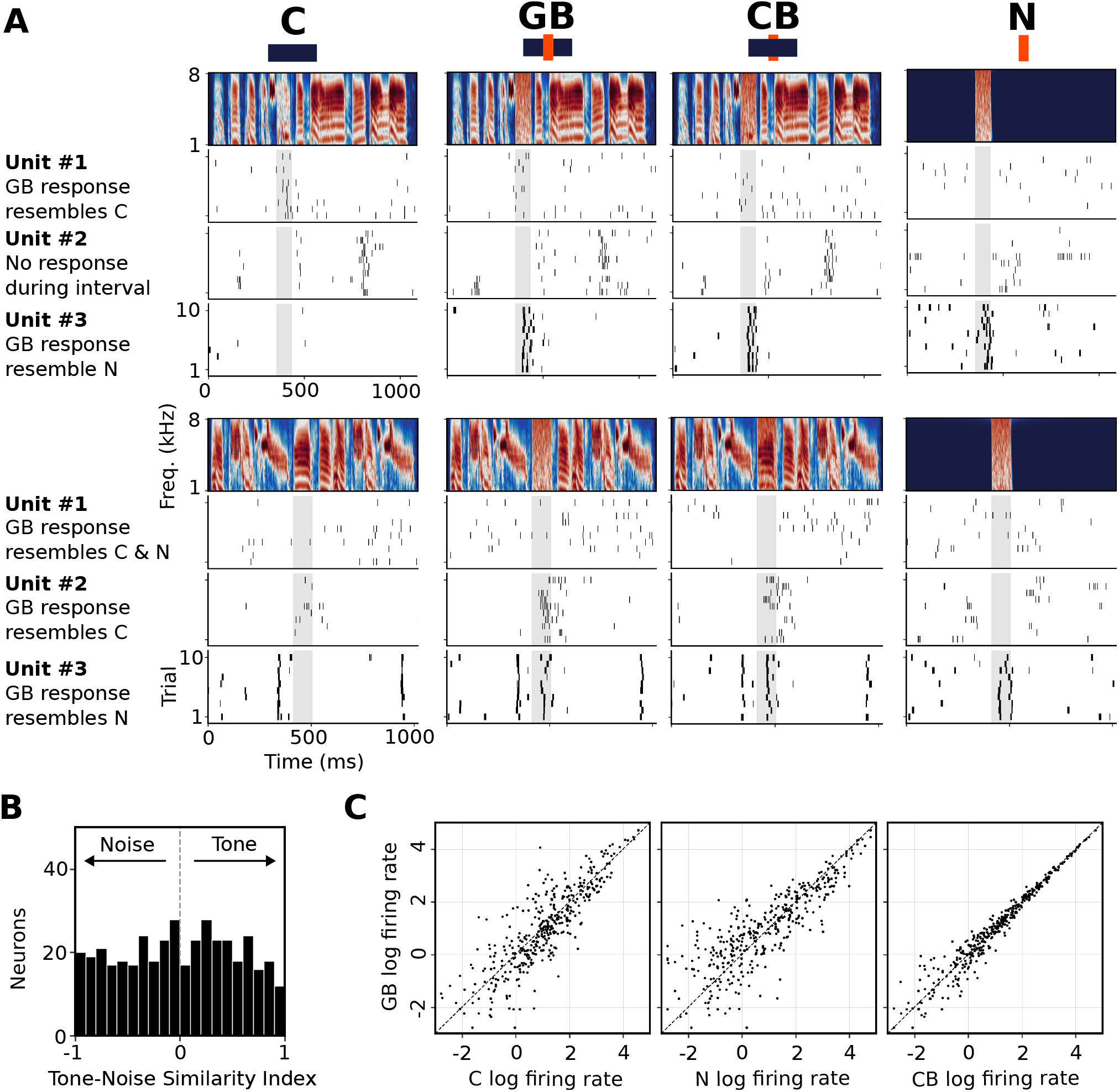
Single unit firing rates during critical intervals. (**A**) Spectrograms of C, GB, CB, and N variants of 2 natural motifs (top and bottom rows), along with raster plots of three single units in response to repeated presentation of stimulus. Shaded region of raster plot indicate critical interval during which motif was modified. Each unit exhibited different firing patterns between variants as well as motifs: unit *−*1 responded more strongly to C, GB, and CBfor top motif but not bottom motif; unit 2 responded to C, GB, and CBfor bottom motif but not top motif; unit 3 responded strongly to GB, CB, and N for both motifs. (**B**) Each recorded single unit’s averaged firing rate during critical intervals of GB stimuli can be measured in similarity to its response to either syllable/tone (C variant) or noise (N variant) using a Tone-Noise Similarity Index (^14^): 1 if GB firing rate is identical to noise, 1 if GB firing rate is identical to tone. Neuron population (*n* = 407 units) showed diverse firing rates during critical interval, TNSI evenly distributed between *−*1 and 1. (**C**) Average firing rates (log scale) during the critical interval of GB for all units, plotted against response rates to C, N, and CB during the same interval. Each point corresponds to an individual unit. Diagonal dotted lines represent equality. CB response rates are highly predictive of GB response rates, while the rates for C and N show more scatter.

**Supplemental Figure 3.**
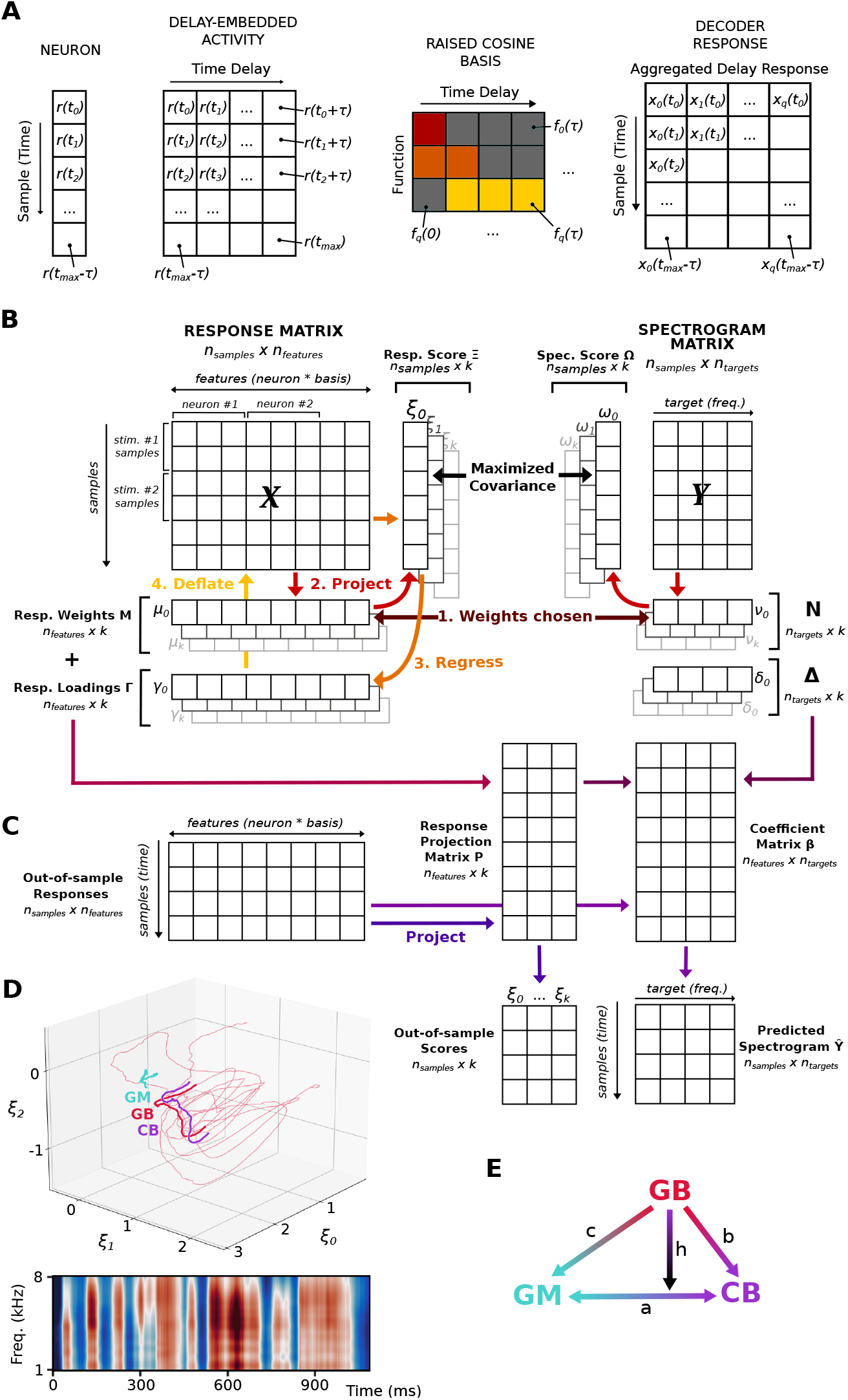
Partial Least Squares decoder. (**A**) Response transformation from average firing rates (peristimulus time histogram), delay-embedding with time window *τ* to form a Hankel matrix, and projection into a raised-cosine basis set. (**B**) Illustration of partial least squares regression (see Methods for details). (**C**) Projection of out-of-sample responses into latent subspace uses matrix **P** calculated from the weight **M** and loading **Γ** matrices. **P** and spectrogram loading matrix **?** can also be used to calculate a coefficient matrix **fi** that produces stimulus spectrogram predictions from responses. (**D**) Top: example projection of neural responses to GM, GB, and CB variants of a motif in the subspace defined by the top three components of the model. Bottom: predicted stimulus spectrogram from GB response. Coefficient matrix calculated from all eleven components of PLS model. (**E**) The Restoration Index (RI) used average distances between the latent neural states of GB, GM, and CB to assess restoration. Geometrical abstract of the RI calculation: given the average Euclidean distances *a, b*, and *c* between pairs of GB, GM, and CB in two or more dimensions, GB is closer to CB if *c − b* is positive. The triangular height h is used to scale the difference: a smaller h means compare GB’s proximity between GM and CB is more relevant.

**Supplemental Figure 4.**
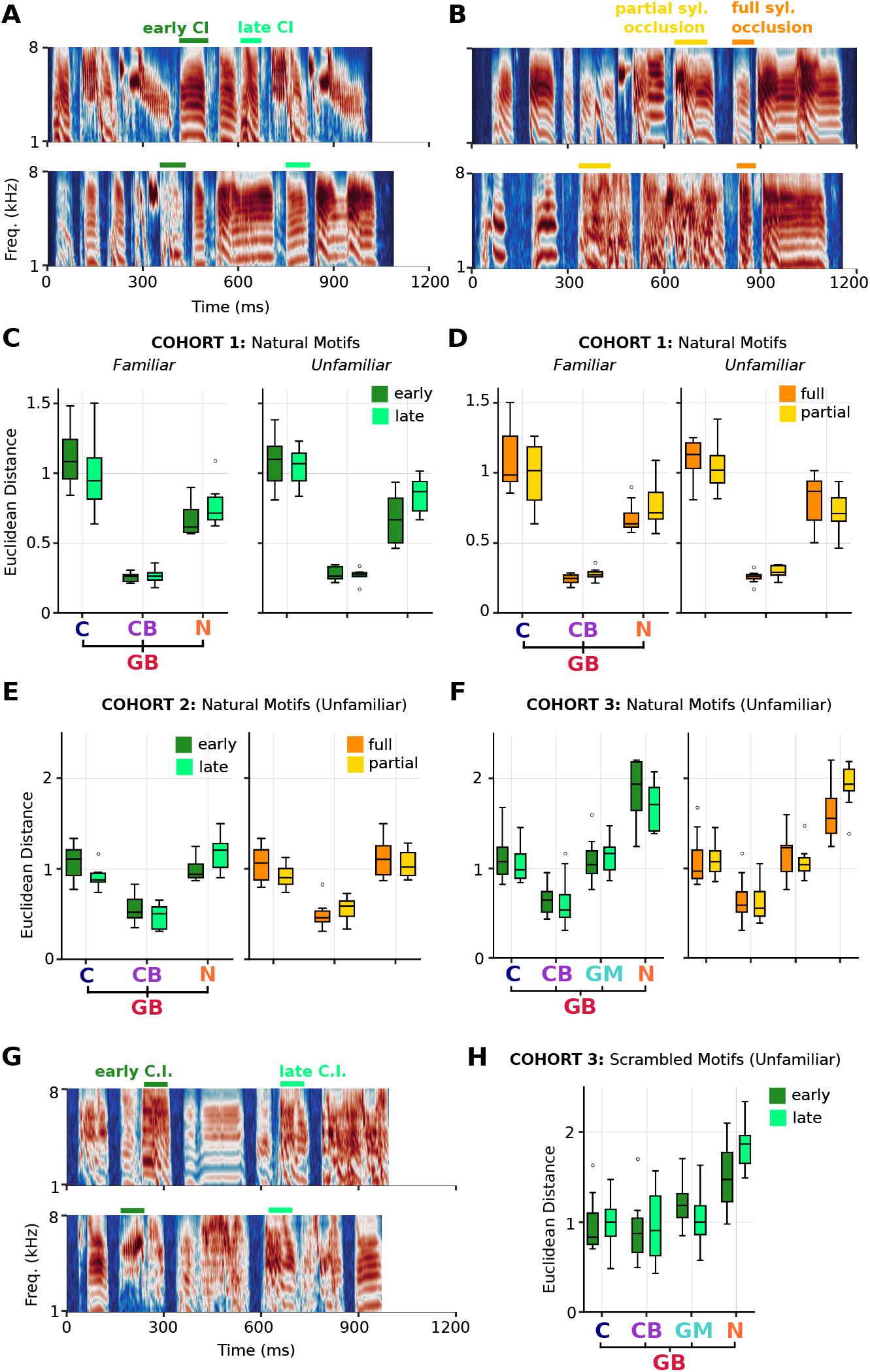
Effects of gap location and occlusion type. (**A**) Stimulus spectrograms of two example natural motifs. Colored bars above indicate location of two critical intervals: one earlier and one later in the motif. (**B**) Spectrograms of two other natural motifs, illustrating partial and full syllable occlusion. (**C**) Decoding results from cohort 1 measured by GB’s Euclidean distance to C, CB, and N, separated by familiarity to motif (left: motifs that subjects were exposed to, right: motifs that subjects had never heard). (**D**) Similarly, no difference found between fully occluded and partially occluded syllables. (**E**) No difference found for projected GB’s distance to C, CB, and N in second cohort of birds, in terms of critical interval location (left) or syllable occlusion type (right). (**F**) Similarly, no difference found in third cohort of birds, where noise-masking stimuli were added. Here, GB’s distance to C, CB, GM, and N were similar regardless of critical interval location or occlusion type. (**G**) Stimulus spectrograms of two example scrambled syntax motifs used. Colored bars above indicate critical interval location: early (second syllable) or late (fourth syllable). All scrambled-syntax motifs’ critical intervals were partial syllable occlusions. (**H**) No difference found in response to scrambled-syntax motif for third bird cohort between early and late critical intervals.

